# Disease-associated mutations in GluK2 disrupt compartment-specific regulatory mechanisms controlling kainate receptor membrane trafficking

**DOI:** 10.64898/2026.05.29.728878

**Authors:** Nora Ibrahim Alorf, Busra P. Yucel, Kevin A. Wilkinson, Jeremy M. Henley

## Abstract

Missense mutations in the *Grik2* gene that encodes the kainate receptor subunit GluK2 are associated with neurodevelopmental disorders, but how individual mutations perturb receptor regulation remains unclear. To investigate how disease-associated GluK2 mutations affect receptor trafficking and signalling we used lentiviral approaches to knockdown and replace endogenous GluK2 with tagged wild-type or mutant GluK2 constructs in primary rat neuronal cultures. We show that two disease-associated GluK2 mutations disrupt receptor trafficking through distinct molecular mechanisms. The neurodevelopmental disorder-associated A657T mutation impairs receptor folding and assembly, resulting in endoplasmic reticulum retention, proteasome-dependent degradation, and severely reduced surface expression. In contrast, the C-terminal mutation M867I, identified in patients with autism, markedly reduces PKC-dependent phosphorylation at S868. This does not impede maturation but uncouples GluK2 surface redistribution from activity-dependent signalling. Thus, pathogenic GluK2 variants selectively disrupt either receptor biogenesis or post-translational regulatory integration. Functionally, these mutations differentially alter downstream Ca²⁺ signalling responses, linking molecular defects to altered receptor output. These findings establish the GluK2 C-terminus as a critical control node for synaptic receptor homeostasis and illustrate how discrete mutations can uncouple distinct layers of membrane protein regulation.

---

Kainate receptors (KARs) are dynamically regulated membrane proteins that assemble as tetramers composed of GluK1-5 subunits, with GluK2/GluK5 heteromers representing the predominant postsynaptic configuration in many brain regions (Petralia *et al*. 1994; Watanabe-Iida *et al*. 2016). KARs contribute to regulating excitatory transmission and synaptic plasticity so their protein folding, receptor assembly, and trafficking are all tightly coordinated to ensure synaptic stability and function (Henley *et al*. 2014; Evans *et al*. 2019; Blakemore *et al*. 2018). Differences in combinatorial assembly with auxiliary proteins, extensive protein-protein interactions, and dynamic post-translational modifications (PTMs) contribute to regulating GluK2-containing KAR surface expression and signalling output (González-González *et al*. 2012; Lerma & Marques 2013; Carta *et al*. 2014; Evans *et al*. 2017).

The intracellular C-terminus of GluK2 integrates multiple regulatory PTM sites, including phosphorylation (Nasu-Nishimura *et al*. 2010), palmitoylation (Pickering *et al*. 1995; Copits & Swanson 2013; Yucel *et al*. 2023), SUMOylation (Martin *et al*. 2007; Konopacki *et al*. 2011; Chamberlain *et al*. 2012), and ubiquitination (Salinas *et al*. 2006; Maraschi *et al*. 2014), which together coordinate receptor surface stability and activity-dependent internalisation (**Figure 1**). While these PTMs are known to regulate mature receptors, it is unclear how disease-associated mutations perturb this regulatory architecture.

**Figure 1.**
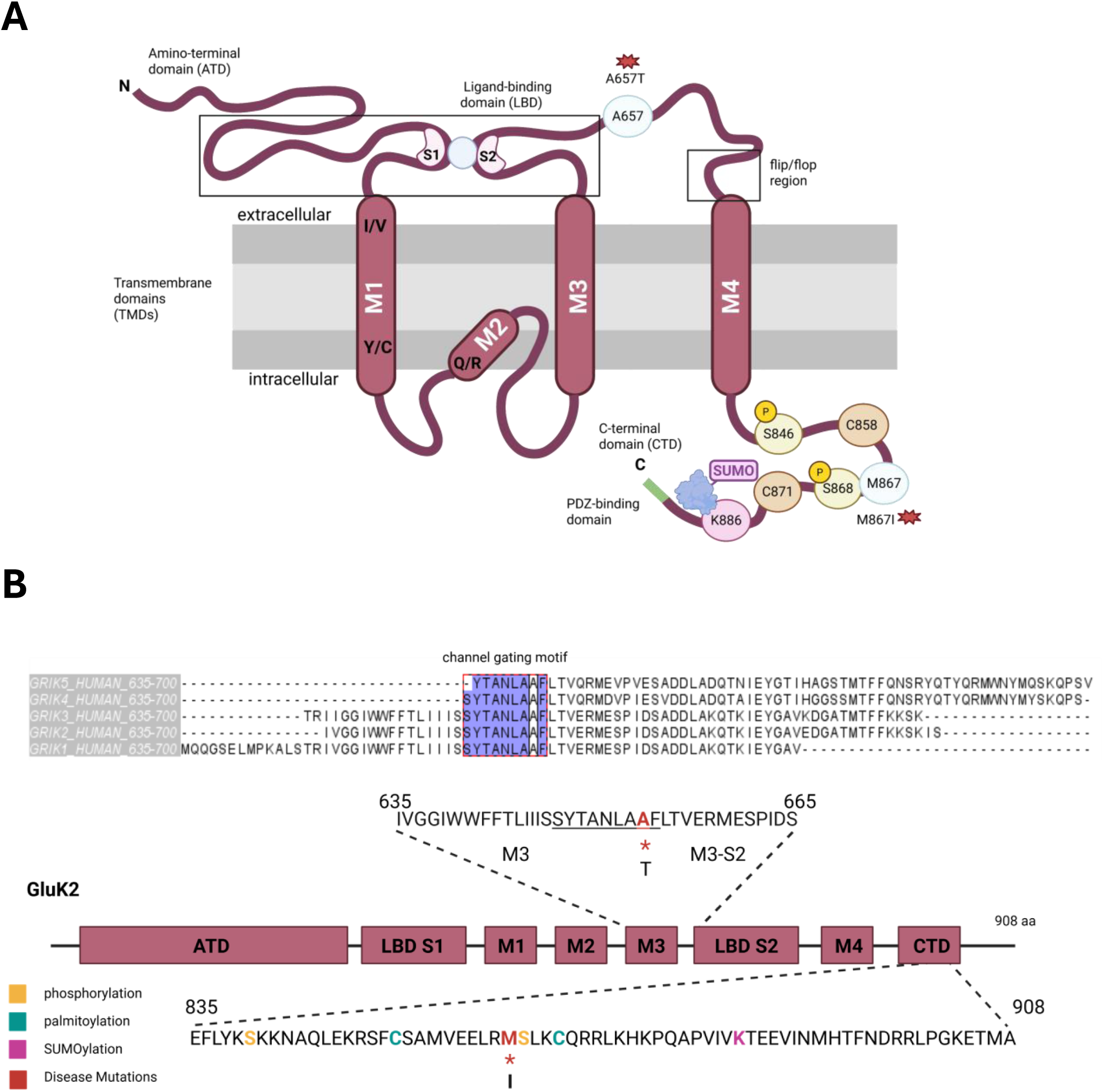
Schematic representations of GluK2 (A) GluK2 subunit topology showing the N-terminal domain (ATD), the ligand-binding domain (LBD; S1 and S2), the transmembrane domains (M1–M4), and the intracellular C-terminal domain (CTD) containing the PDZ-binding motif and key PTM sites. The locations of the disease-associated mutations A657T (within the M3 pore-forming region) and M867I (within the CTD, proximal to the PKC phosphorylation site) are indicated by red asterisks. (B) Linear schematic of GluK2 showing domain organisation and key sequence features, with the position of the M867I mutation indicated within the C-terminal region (bottom). Top, sequence alignment of the M3 channel-gating motif (SYTANLAA) across kainate receptor subunits, highlighting the position of A657 (asterisk) affected by the A657T mutation.

Missense variants in the *Grik2* gene that encodes GluK2 have been identified in patients with intellectual disability and epilepsy (Matute 2011; Fritsch *et al*. 2014; Henley *et al*. 2014; Barthet *et al*. 2022; Chalupnik & Szymanska 2023), yet the molecular consequences of individual mutations remain poorly defined.

We investigated two pathogenic GluK2 mutations in different structural domains. The C-terminal mutation M867I, identified in individuals with autism spectrum disorder (Jamain *et al*. 2002), has previously been described as a gain-of-function variant associated with enhanced plasma membrane expression and increased kainate-evoked currents in *Xenopus* oocytes (Strutz-Seebohm *et al*. 2006). This increase has been proposed to arise from the location of the mutation near a trafficking motif (Yan *et al*. 2004) (Han *et al*. 2010). M867I is positioned adjacent to S868, a key PKC phosphorylation site in the GluK2 C-terminus, raising the possibility that this mutation perturbs PTM-dependent regulation of KAR trafficking and surface expression. Consistent with this idea, we found that expression of GluK2(Q)-M867I preserves receptor assembly but perturbs PKC-dependent phosphorylation, resulting in defects in downstream trafficking events. In addition, electrophysiological analyses indicate that GluK2 M867I impair channel gating and slow receptor desensitisation (Han *et al*. 2010).

The A657T mutation within the pore-loop region of GluK2 was originally identified in a patient with severe neurodevelopmental delay and cognitive impairment (Guzman *et al*. 2017) and has been linked to profound defects in channel gating and receptor surface trafficking (Guzman *et al*. 2017; Stolz *et al*. 2021). More recently, analysis of a knock-in mouse model showed that this mutation perturbs neuronal excitability (Nomura *et al*. 2023). Consistent with this, our results show that A657T also disrupts receptor maturation, leading to ER retention and proteasome-dependent degradation.

Here, we reveal mechanistically separable layers of receptor control and demonstrate that disease-associated mutations can perturb either early biogenesis or post-translational regulatory integration. This study further establishes the GluK2 C-terminus as a modular signalling hub and provides a framework for understanding mutation-specific disruption of membrane protein homeostasis.

## Results

### GluK2-A657T primarily disrupts receptor biogenesis

To investigate how disease-associated GluK2 mutations affect receptor surface expression under basal conditions, we used lentiviruses to knockdown (KD) endogenous GluK2 and replace it with a GFP-Myc-tagged GluK2-WT, –M867I, or –A657T equivalent in cortical neurons. Surface expression was quantified using surface biotinylation assays followed by immunoblotting. Under basal conditions, GluK2-A657T exhibited significantly lower surface expression, whereas GluK2-M867I surface levels were comparable to GluK2-WT (**Figure 2A**).

**Figure 2.**
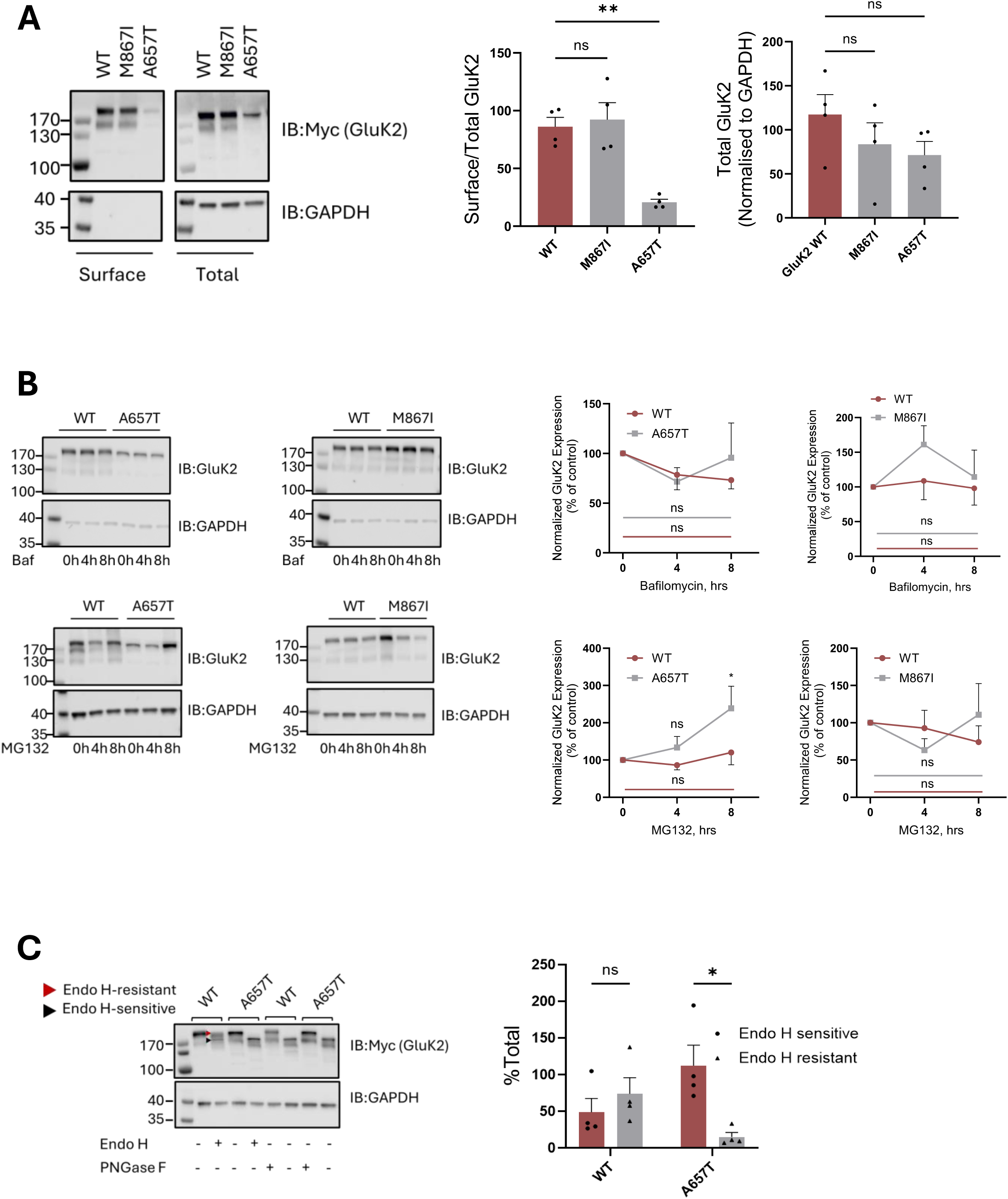
GluK2 A657T causes ER retention and reduced surface trafficking. (A) Representative surface biotinylation immunoblots of cortical neurons expressing GFP-Myc-GluK2 constructs. Graphs show quantification of GluK2 surface levels normalised to total GluK2, with total GluK2 quantified relative to GAPDH. One-way ANOVA with Tukey’s multiple-comparisons test (**p<0.01, ***p<0.001). N=4 independent dissections. (B) Representative immunoblots of cortical neurons infected with KD-replacement viruses expressing GFP-Myc-GluK2-WT, M867I or A657T. Seven days post-infection, cells were treated with 20µM bafilomycin or 20µM MG132, lysed and immunoblotted to assess total GluK2 accumulation over time. Quantification of GluK2 levels normalised to GAPDH. N=4 independent experiments using cells from separate dissections. Two-way ANOVA with Tukey’s multiple-comparisons test (*p < 0.05). (C) Cortical neurons expressing GFP-Myc-GluK2 WT or A657T were lysed, and 5µg of protein was treated with Endo H or PNGase F. Representative anti-Myc immunoblots show GluK2 glycoforms: the upper band corresponds to complex-glycosylated Endo H-resistant, the lower band is Endo H-sensitive (ER-retained) GluK2. The graph shows quantification of Endo H-sensitive and –resistant relative to total GluK2. N=4 independent experiments using cells from separate dissections. Two-way ANOVA with Tukey’s multiple-comparisons test (*p<0.05). All error bars indicate SEM.

To determine whether lower levels of GluK2-A657T result from enhanced protein degradation, we examined the effects of proteasomal and lysosomal pathway inhibition. Total levels of GluK2-WT and GluK2-M867I were similar, but there was increased accumulation of GluK2-A657T following proteasome inhibition with MG132, but not lysosomal inhibition with bafilomycin (**Figure 2B**). These results suggest rapid degradation of misfolded receptors rather than altered synthesis or degradation after endocytosis.

To examine whether GluK2-A657T was misfolded and ER-retained, we next performed glycosylation analysis with Endo H and PNGase F to monitor GluK2 maturation. GluK2-WT exhibited both an Endo H-sensitive, immature form corresponding to the ER-retained receptor and an Endo H-resistant, PNGase F-sensitive form consistent with Golgi-processed mature receptor. In contrast, GluK2-A657T was predominantly Endo H-sensitive, lacking the mature, Golgi-processed form (**Figure 2C**). These data are consistent with GluK2-A657T disrupting the early folding or assembly required for efficient ER exit. Taken together, these results indicate that GluK2-A657T causes a biogenesis defect characterised by impaired maturation and rapid proteasomal degradation, thereby severely limiting the pool of GluK2 available for surface expression. The GluK2-M867I mutation, on the other hand, maintains normal biogenesis and basal trafficking.

### GluK2-M867I selectively disrupts PKC-dependent regulation

Because the M867I mutation lies immediately adjacent to a known PKC phosphorylation site, we next examined PKC-dependent phosphorylation and trafficking of GluK2-M867I and GluK2-A657T. Phos-tag analysis revealed robust PMA-induced phosphorylation of S868 in GluK2-WT, which was not present in the non-phosphorylatable S868A mutant. Interestingly, phosphorylation of GluK2-M867I was also completely abolished, but significantly enhanced in GluK2-A657T under the same conditions (**Figure 3A**).

**Figure 3.**
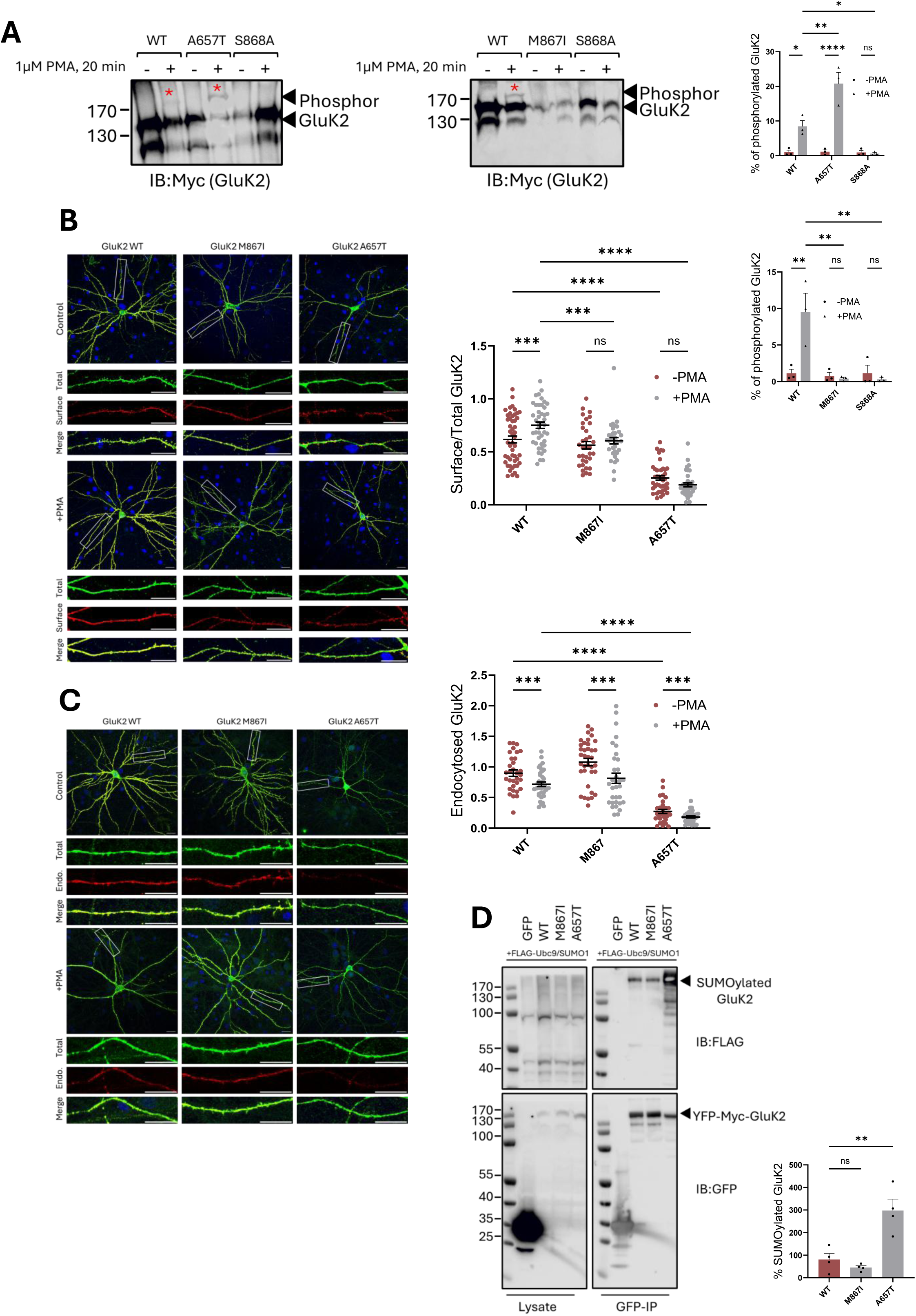
GluK2 M867I disrupts PKC-driven GluK2 trafficking. (A) Seven days post-transduction cortical neurons expressing GFP-Myc-GluK2 constructs were stimulated with 1µM PMA for 20 min and analysed by Phos-tag gel electrophoresis. Representative immunoblots show unphosphorylated and Phos-tag–shifted GluK2 bands and the percentage of phosphorylated GluK2 quantified. N=3 independent experiments using cells from separate dissections. Two-way ANOVA with Tukey’s multiple-comparisons test (*p<0.05, **p<0.01, ****p<0.0001). (B) Representative confocal images of hippocampal neurons transfected with GFP-Myc-GluK2 constructs were stimulated with 1µM PMA for 20min and surface expressed GluK2 was imaged using an anti-GFP antibody (red; Cy3-conjugated secondary). Cells were then fixed with 4% PFA, permeabilised, and stained for total GluK2 (green; anti-GFP antibody with Alexa Fluor 488–conjugated secondary). DAPI (blue), surface (red), total (green), and merged (yellow) GluK2. ROIs are indicated by white boxes, scale bars = 20 µm. Quantification of surface GluK2 levels normalised to total GluK2. N=34-44 cells per condition from 3 independent dissections. Two-way ANOVA with Tukey’s multiple-comparisons test (***p<0.001, ****p<0.0001). (**C**) Antibody-feeding assay of GFP-Myc-GluK2 internalisation in hippocampal neurons. Seven days post-infection, live cells were incubated with an anti-GFP antibody (45 min), followed by PMA stimulation (1 µM, 20 min). Surface-bound antibodies were removed by acid stripping and cells were fixed with 4% PFA. Total GluK2 was detected using an anti-GFP antibody with Alexa Fluor 488–conjugated secondary antibody (green), and internalised GluK2 was detected using a Cy3-conjugated secondary antibody (red). Scale bar in ROI = 20 µm. Quantification of internalised GFP-Myc-GluK2 normalised to total GFP-Myc-GluK2 expression. N=32-36 cells per condition from 3 independent dissections. Two-way ANOVA with Tukey’s multiple-comparisons test (***p<0.001, ****p<0.0001). (**D**) HEK293T cells were transfected with a GFP-Myc-tagged GluK2 construct, FLAG-Ubc9 and FLAG-SUMO1. 48–72h post-transfection, GFP-Myc-tagged GluK2 was isolated using GFP trap beads and immunoblotted. SUMOylated GluK2 was detected with anti-FLAG antibody. SUMOylation was normalised to the corresponding total GFP-Myc-GluK2 construct expression. N=4 independent experiments. One-way ANOVA with Tukey’s multiple-comparisons test (***p<0.001) All error bars indicate SEM.

We next measured the response of WT or mutant GluK2 to PKC activation. We have shown previously that activation of PKC with PMA leads to an increase in surface expression of GluK2 (Chamberlain *et al*. 2012). As expected, activation of PKC with PMA (1μM, 20 min) significantly increased surface expression of GluK2-WT but failed to enhance the surface to total ratio of either GluK2-M867I or GluK2-A657T (**Figure 3B**).

We have shown previously that, as well as enhancing GluK2 surface expression through enhanced receptor recycling, activation of PKC also leads to enhanced rates of receptor endocytosis by promoting GluK2 SUMOylation at K886 (Konopacki *et al*. 2011; Chamberlain *et al*. 2012). Therefore, we next measured endocytosis rates of GluK2 in hippocampal neurons to determine whether the impaired PKC responsiveness of GluK2-M867I reflects altered receptor internalisation. Cell surface labelling to follow endocytosis confirmed that GluK2-M867I internalisation was comparable to GluK2-WT, indicating that the mutation does not affect PKC-dependent internalisation, but more likely disrupts PKC-dependent surface delivery. (**Figure 3C**). Surface levels of GluK2-A657T were very low under basal conditions, which we attribute to enhanced proteasomal degradation before the subunit reaches the plasma membrane. Nonetheless, PMA further reduced GluK2-A657T endocytosis. Phosphorylation of GluK2 at S868 enhances GluK2 SUMOylation at K886 (Konopacki *et al*. 2011), so we next examined SUMOylation levels of our GluK2 mutants. Under basal conditions, SUMOylation of GluK2-A657T was enhanced compared to GluK2-WT (**Figure 3D**). SUMOylation of GluK2-M867I was not significantly altered compared to GluK2-WT (**Figure 3D**). These results suggest that M867I does not affect GluK2 basal surface expression or maturation, but rather it disrupts PKC-dependent surface trafficking.

### Post-translational modifications underpin differential trafficking

The non-SUMOylatable mutant GluK2-K886R displays higher levels of basal surface expression (Martin *et al*. 2007), consistent with preventing SUMOylation at K886, reducing internalisation. GluK2-M867I displayed reduced S868 phosphorylation (**Figure 3A**), but basal surface delivery was unaffected.

Further characterisation of post-translational modifications of GluK2 revealed complementary mechanisms underpinning the observed trafficking phenotypes. GluK2 is palmitoylated at two cysteine residues, C858 and C871, which stabilise receptors at the plasma membrane in an activity-dependent manner (Yucel *et al*. 2023). GluK2-A657T showed significantly reduced palmitoylation (**Figure 4A**), compared to GluK2-M867I mutant, which also showed moderately reduced palmitoylation compared to GluK2-WT. Together with increased SUMOylation and phosphorylation of GluK2-A657T, these results are consistent with rapid internalisation of the small fraction of GluK2-A657T that reaches the plasma membrane.

**Figure 4.**
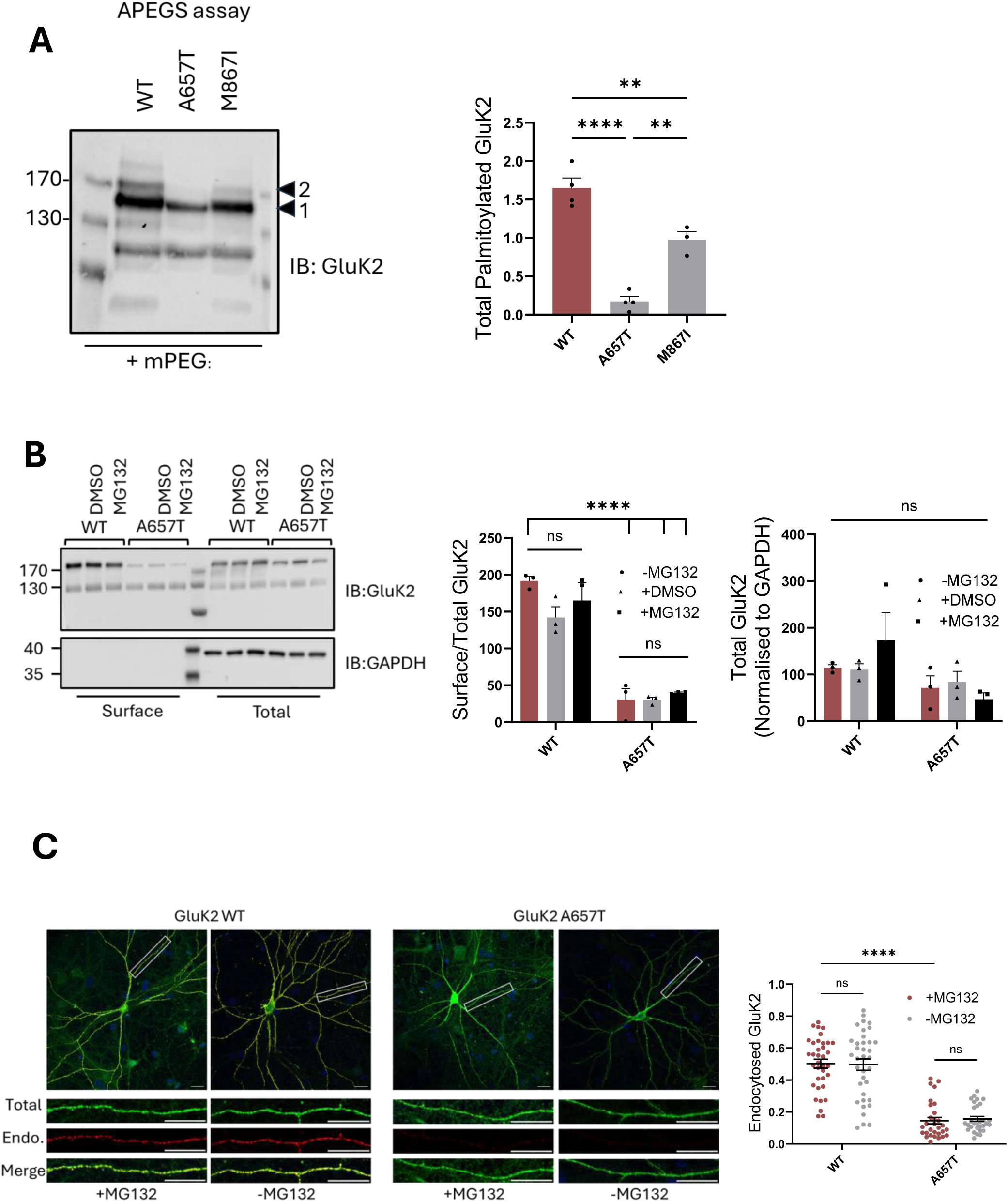
Imbalance of post-translational modifications contributes to A657T instability (A) Seven days after transfection cortical neurons expressing GFP-Myc-GluK2 (WT, A657T, and M867I) constructs were subjected to APEGS assay. Representative blots show upper and lower bands representing single and double palmitoylation. One-way ANOVA was used, followed by Tukey’s multiple-comparisons test (**p<0.01, ****p<0.0001). N=3 independent experiments using cells from different dissections. (B) Representative immunoblots of cortical neurons infected with KD replacement viruses expressing GFP-Myc-GluK2 WT or A657T. Seven days post-infection, cultures were treated with 20µM MG132 or DMSO vehicle for 8h, surface proteins biotinylated and blotted. GAPDH was used as a loading control to verify that intracellular proteins were not biotinylated. Quantification of surface GluK2 normalised to total GluK2 and total GluK2 to GAPDH. N=3 independent dissections. Two-way ANOVA with Tukey’s multiple-comparisons test (***p<0.0001). (C) Hippocampal neurons transfected with GFP-Myc–GluK2 WT or A657T. Seven days post-transfection, cells were treated with 20μM MG132 or DMSO for 4h. Cells were live labelled with an anti-GFP antibody for 45 min, acid washed to remove uncoupled antibody, fixed, permeabilised, and internalised receptors detected with Cy3-secondary antibody (red). Total GluK2 was labelled with anti-GFP and 488 secondary antibody (green); nuclei were stained with DAPI (blue). Scale bars = 20 µm. N=30–36 cells per condition from 3 independent dissections. Statistical analysis was performed using two-way ANOVA with Tukey’s multiple-comparisons test. ****p<0.0001. Error bars indicate SEM.

We next showed that increasing total levels of GluK2-A657T by preventing proteasomal degradation using MG132 does not enhance its traffic to the surface. These data indicate that a trafficking defect leading to ER retention rather than increased protein turnover underlies poor surface expression of GluK2-A657T (**Figure 4B-C**).

Since only a small proportion of GluK2-A657T receptors reach the surface, we propose that the reduced palmitoylation of those receptors could contribute to their rapid removal from the membrane through accelerated internalisation. Alternatively, palmitoylation may occur only at surface-expressed receptors, so the reduced level of palmitoylation may be attributable to fewer receptors trafficking to the plasma membrane. Thus, it remains unclear whether reduced palmitoylation is a cause or consequence of impaired trafficking.

Overall, our data indicate that the integrated actions of phosphorylation, SUMOylation and palmitoylation provide the molecular basis for rapid turnover and limited surface expression of GluK2-A657T, whereas GluK2-M867I primarily affects responsiveness regulatory mechanisms.

### Functional Consequences of Mutations

Pre-mRNA Q/R editing of the *Grik2* gene changes a genetically encoded glutamine (Q) in the channel pore region to an arginine (R) in the GluK2 protein. The unedited GluK2(Q) more efficiently exits the ER, is better surface expressed, and KARs containing GluK2(Q) are Ca^2+^-permeable (Gurung *et al*. 2018; Evans *et al*. 2019; Nair *et al*. 2023).

We exploited these properties of GluK2(Q) for live-cell calcium imaging in HEK293T cells expressing the Ca^2+^-indicator R-GECO to assess how GluK2-A657T and GluK2-M867I affect KAR function. We used the Ca^2+^-impermeable GluK2(R)-WT subunit as a control (Gurung *et al*. 2018; Evans *et al*. 2019; Nair *et al*. 2023). Both GluK2 disease mutants remain responsive to kainate, but display altered signalling compared to GluK2(Q)-WT. GluK2-M867I exhibited reduced response amplitude but achieved endpoints similar to GluK2(Q)-WT, whereas GluK2-A657T showed reduced sustained signalling and altered kinetics (**Figure 5**). These data indicate that GluK2-A657T not only limits surface receptor availability but also alters functional Ca²⁺ responses.

**Figure 5.**
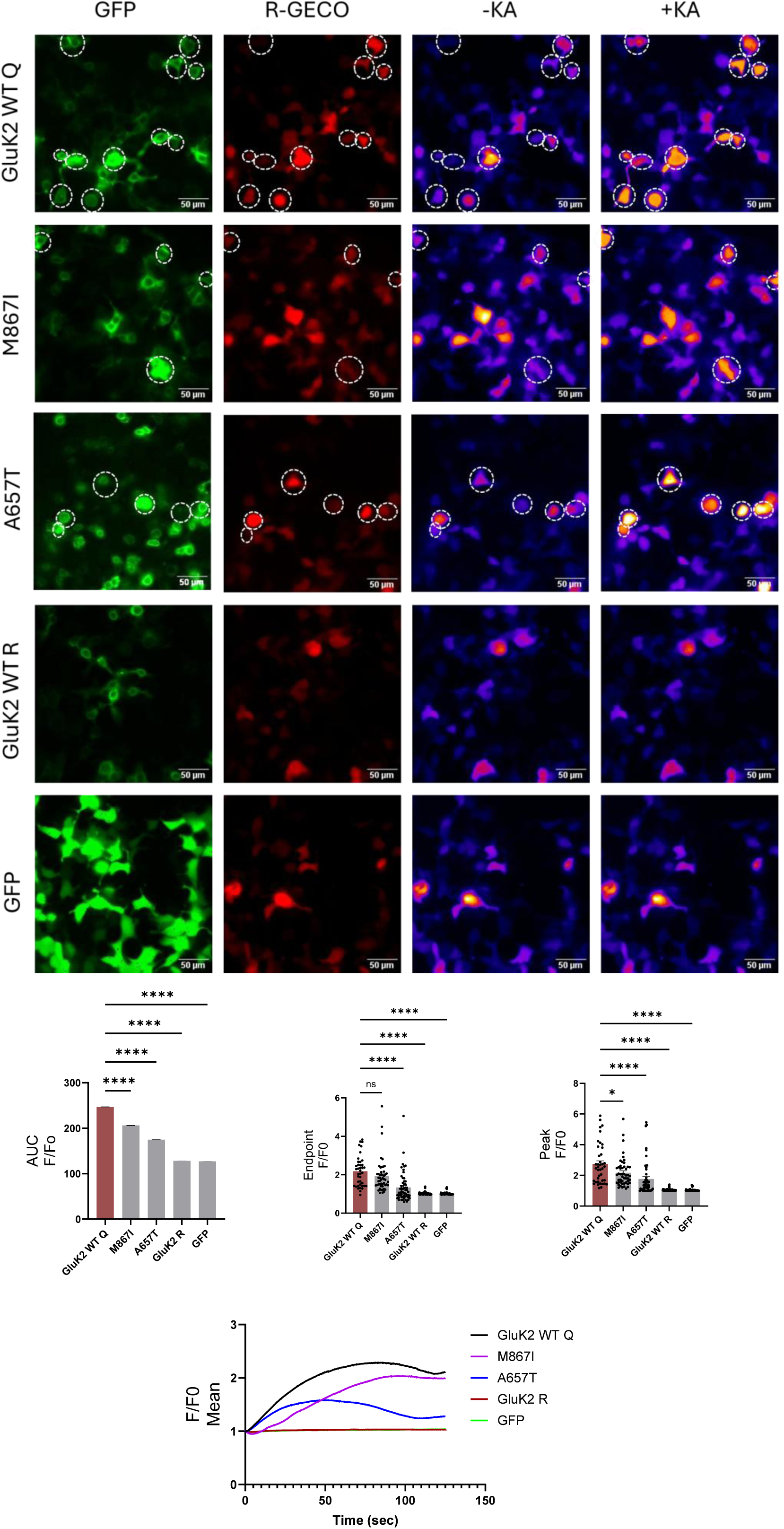
Calcium imaging of GluK2 mutant function reveals altered amplitude and kinetics. HEK293T cells were transfected with GFP or GFP-Myc-tagged GluK2 constructs (WT Q, WT R, M867I or A657T) together with the calcium indicator R-GECO. At 48 h post-transfection, cells were stimulated with 100µM KA and calcium activity was recorded for 2 min using widefield imaging. Representative images show GluK2/GFP expression (green) and R-GECO signal (red). Dashed circles indicate ROIs used for quantification. Pseudocolour images illustrate the R-GECO response before (−KA) and after (+KA) KA application. Graphs show fluorescence traces (F/F₀) over time for each condition, aligned to KA application. Scatter plots show endpoint F/F₀, peak F/F₀, and area under the curve (AUC) for the R-GECO response. GluK2 mutations alter kainate-evoked Ca²⁺ signalling. N=40-60 cells from 3 independent experiments. Statistical analysis was performed using one-way ANOVA with Tukey’s multiple-comparisons test (*p<0.05, ****p<0.0001). Error bars indicate SEM.

## Discussion

In this study, we demonstrate that two disease-associated GluK2 mutations disrupt kainate receptor regulation through fundamentally distinct mechanisms: one targeting receptor biogenesis and the other selectively impairing activity-dependent control. These findings refine our understanding of how pathogenic variants in *GRIK2* perturb synaptic receptor homeostasis and further highlight the GluK2 C-terminus as a vulnerable integration hub for post-translational regulation.

The A657T mutation in the *GRIK2* gene, encoding the GluK2 kainate receptor subunit, has been associated with neurodevelopmental disorders. It occurs within the M3 transmembrane helix, a critical structural element forming the channel gate of ionotropic glutamate receptors (Guzman *et al*. 2017). Previous electrophysiological studies demonstrated that the A657T mutation in GluK2 produces KARs with slowed deactivation and reduced desensitisation. These changes prolonged channel opening and consequent ion flux resulting in a gain-of-function phenotype (Stolz *et al*. 2021). Consistent with these functional alterations, knock-in mouse models show increased neuronal excitability and synaptic dysfunction (Nomura *et al*. 2023).

We show that in addition to altered channel kinetics, the A657T mutation profoundly disrupts GluK2-containing KAR trafficking and post-translational regulation. GluK2-A657T subunits are predominantly Endo H–sensitive, accumulate upon proteasome inhibition, and have dramatically reduced plasma membrane expression. These findings indicate that GluK2-A657T is regulated by ER quality-control pathways and ER-associated degradation.

Moreover, our data are also consistent with the small population of GluK2-A657T containing KARs that reach the plasma membrane being rapidly destabilised and internalised due to reduced palmitoylation and enhanced phosphorylation and SUMOylation. These data agree with the recent proposal that coordinated interplay between these PTMs promotes GluK2-containing KAR internalisation (Yucel *et al*. 2023).

KARs containing the edited GluK2(Q) subunit are permeable to Ca²⁺ and have previously been used to monitor receptor activity through agonist-evoked calcium signals in heterologous systems (Burnashev *et al*. 1995; Lerma & Marques 2013). We show that kainate application induces robust Ca²⁺ responses in cells expressing GluK2(Q)-WT, whereas the Ca²⁺-impermeable GluK2(R)-WT control elicited only a minimal signal (**Figure 5**). Although both GluK2(Q)-A657T and GluK2(Q)-M867I were responsive to kainate, they exhibited different Ca²⁺ signalling profiles. GluK2(Q)-A657T displayed reduced sustained signalling and altered kinetics, consistent with our trafficking data showing markedly reduced surface receptor expression. These findings suggest that, beyond previously described gating defects of GluK2-A657T (Guzman *et al*. 2017; Stolz *et al*. 2021), this mutation also limits functional receptor availability at the plasma membrane, thereby reshaping downstream Ca²⁺ signalling. Together, these findings indicate that the A657T mutation disrupts GluK2 function through multiple mechanisms, including impaired receptor trafficking, altered post-translational regulation, and modified Ca²⁺ signalling, suggesting that the pathogenic effects of this variant arise from a combined defect in receptor surface availability and channel function.

In contrast to GluK2-A657T, the GluK2-M867I mutation does not impair receptor maturation or basal surface delivery, indicating that receptor biogenesis and constitutive trafficking remain largely intact. Previous work characterising this mutant reported only modest effects on channel kinetics, including a slight slowing of desensitisation, suggesting that GluK2-M867I does not strongly disrupt receptor gating (Han *et al*. 2010). However, the proximity of M867 to the PKC-regulated S868 phosphorylation site raised the possibility that this region may play a role in receptor regulation. The intracellular C-terminus of GluK2 integrates phosphorylation (Nasu-Nishimura *et al*. 2010), SUMOylation (Martin *et al*. 2007) and palmitoylation (Yucel *et al*. 2023) to coordinate receptor trafficking and turnover.

The M867I mutation reduces PKC phosphorylation and blocks PKC-dependent recycling/exocytosis. Surprisingly, however, PKC-dependent endocytosis and SUMOylation remain intact. Thus, this mutation does not compromise receptor structural integrity but instead disrupts dynamic post-translational regulatory control.

These mechanistically divergent disruptions resulting from the A657T and M867I point mutations in GluK2 are likely to have distinct consequences at the circuit level. The impaired receptor biogenesis observed for GluK2-A657T would reduce the availability of functional KARs during critical periods of synaptic development, potentially compromising excitatory-inhibitory balance and network maturation. In contrast, the GluK2-M867I retains basal expression but disrupts activity-dependent redistribution, which may impair synaptic plasticity and dynamic tuning of neuronal excitability (**Figure 6**).

**Figure 6.**
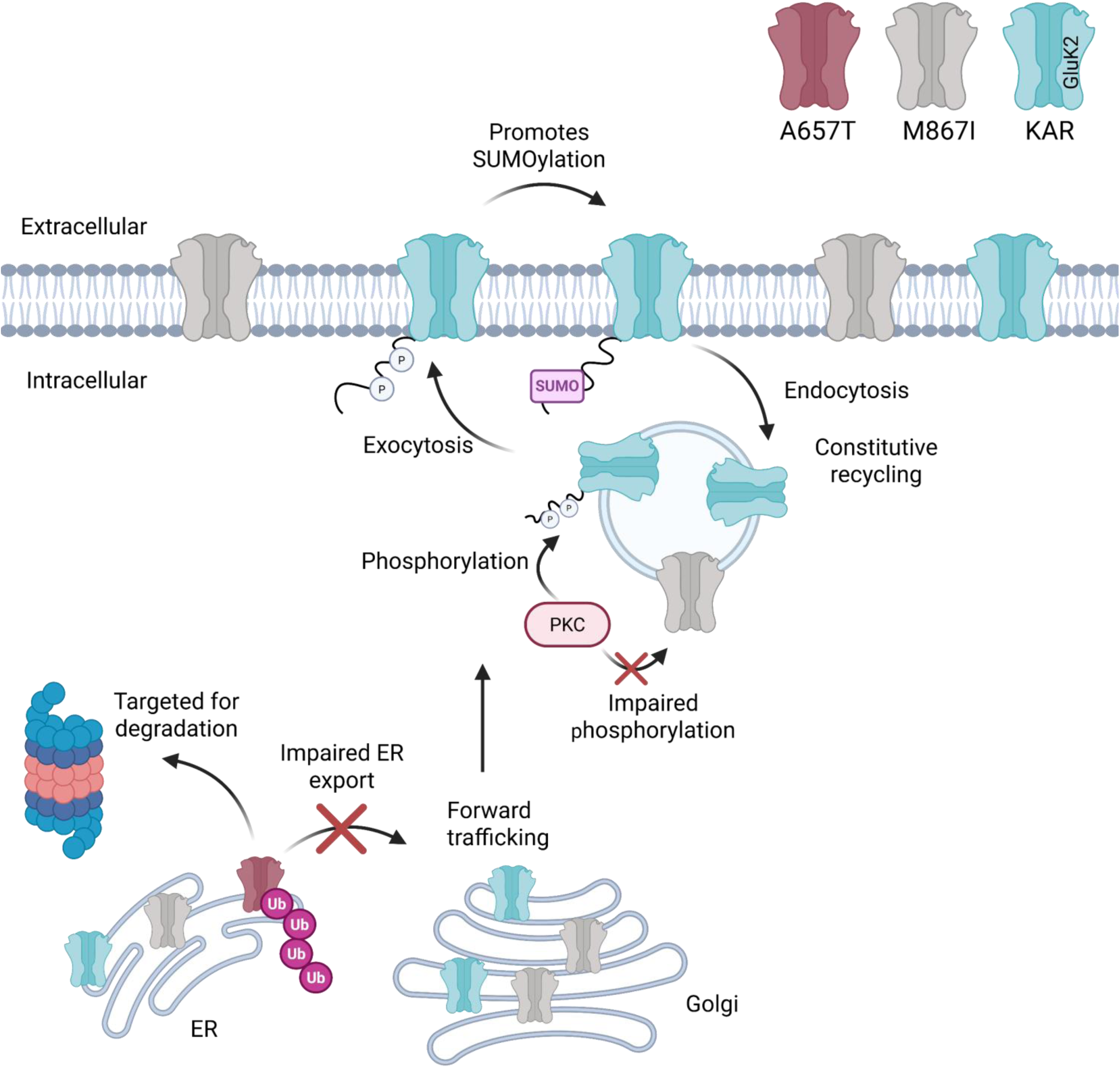
Schematic model of the effects of disease-linked GluK2 mutations on receptor trafficking. GluK2 A657T displays a trafficking defect within the secretory pathway and fails to efficiently exit the ER, likely owing to impaired folding or disrupted subunit assembly. Consequently, the mutant is targeted for degradation, resulting in reduced surface expression. By contrast, GluK2 M867I reaches the cell surface under basal conditions but disrupts activity-dependent PKC-mediated trafficking. Under normal conditions, PKC-dependent phosphorylation of GluK2 enhances receptor trafficking at the plasma membrane, which in turn promotes SUMOylation and subsequent receptor internalisation.

Overall, our findings identify two mechanistic classes of pathogenic GluK2 mutation: defects in biogenesis and defects in regulatory control of receptor trafficking. These results further highlight the GluK2 C-terminal domain as a critical control hub for PTM-dependent receptor trafficking and demonstrate how discrete perturbations within a single receptor subunit can disrupt synaptic homeostasis at multiple levels. More broadly, our study illustrates how discrete mutations within a single membrane protein can uncouple distinct regulatory modules, providing a framework for understanding mechanistic diversity in protein mis-regulation. Future studies will be required to determine how these distinct molecular mechanisms influence synaptic function and circuit activity in neurons and *in vivo*.

## Material and methods

### Primary antibodies

| Antibody | Supplier | Catalog number | Species | Dilution |
| --- | --- | --- | --- | --- |
| GluK2 (WB) | Millipore | 04-921 | Rabbit | 1:5000 |
| GFP (WB) | ChromoTek | 3H9-100 | Rat | 1:5000 |
| GFP (ICC) | Abcam | ab13970 | Chicken | 1:1000 |
| Myc (WB) | Cell Signalling | 71D10 | Rabbit | 1:1000 |
| FLAG (WB) | Sigma | F3165 | Mouse | 1:1000 |
| GAPDH (WB) | Abcam | ab8245 | Mouse | 1:10000 |

### Primary neuronal culture

All the animal experiments and procedures were performed in compliance with the UK Animal Scientific Procedures act (1986) and were guided by the Home Office Licensing Team at the University of Bristol. All animal procedures relating to this study were approved by the Animal Welfare and Ethics Review Board at the University of Bristol (approval number UIN/18/004).

Primary hippocampal and cortical cultures were prepared from embryonic day 18 (E18) Wistar rat embryos as described previously (Nair et al., 2021). Briefly, embryos were removed into Hanks’ Balanced Salt Solution (HBSS), and hippocampal and cortical tissues were dissected, washed, and incubated in 0.005% trypsin/EDTA at 37 °C (10 min for hippocampus, 15 min for cortex). After washing, tissues were triturated in plating medium (Neurobasal supplemented with 5% horse serum, 1% penicillin–streptomycin, 1% GlutaMAX, and 2% B27), filtered, and cells counted. Cells were plated on poly-L-lysine–coated plates at 500,000 cells per well for biochemical assays or 200,000 cells per well for immunocytochemistry. After 2 h, plating medium was replaced with feeding medium (Neurobasal supplemented with 1% penicillin–streptomycin, 1% GlutaMAX, and 2% B27), and cultures were supplemented with additional feeding medium after 7 days in vitro.

### HEK293T cells transfection

Human embryonic kidney (HEK) 293T cells (ECACC) were cultured in Dulbecco’s modified Eagle’s medium (DMEM) containing L-glutamine, supplemented with 10% fetal bovine serum (FBS) and penicillin–streptomycin (Gibco) (complete DMEM), at 37°C in 5% CO₂. Cells were passaged every 3 to 4 days at 80-90% confluency. For passaging, cells were washed with 1xDulbecco’s phosphate-buffered saline (PBS; Thermo Fisher) and detached using 0.05% trypsin (Gibco).

HEK293 cells were transfected 24h after plating or used for lentiviral production. For transfection, plasmid DNA encoding the proteins of interest was diluted in plain DMEM, and Lipofectamine (Invitrogen, 1.5µl per µg DNA) was added. After a 30 min incubation at room temperature to allow complex formation, cells were washed with pre-warmed DMEM supplemented only with 10% FBS and then the transfection mixture was added. Cells were incubated at 37°C in 5% CO₂, and experiments were performed 48h-72h post-transfection.

### Lentivirus preparation

Lentivirus were prepared as described (Wilkinson *et al*. 2022) using polyethylenimine (PEI). Briefly, 4µg plasmids encoding the XLG shRNA transfer vector, 1µg pMD2.G and 3µg p8.9 packaging plasmid were diluted in 1ml plain DMEM. In parallel, PEI (24µL of 1mg/ml) was diluted in 1ml plain DMEM, incubated for 2-3 min at room temperature, then combined with the DNA mixture and incubated for 30 min at room temperature to allow complex formation. HEK293T cells were washed once with pre-warmed DMEM and incubated with the transfection mixture for 4h at 37°C and 5% CO₂. The transfection medium was then replaced with complete DMEM, and cells were returned to the incubator for 48h. Viruses were harvested 48h post-transfection, centrifuged to remove cellular debris, and passed through a 0.45µm filter. The supernatant was aliquoted and stored at −80°C until use. Primary cortical neurons were transduced at DIV12-14 and maintained for 7 days to allow sufficient expression/KD prior to experiments.

### Surface biotinylation

Surface proteins were labeled with Sulfo-NHS-SS-Biotin (Thermo Fisher, Cat# 21331) and subsequently pulled down using streptavidin beads (Merck, Cat# S1638), as previously described (Nair *et al*. 2021). Samples were subjected to SDS–PAGE and Western blotting, and membranes were probed with anti-Myc, anti-GluK2 and anti-GAPDH antibodies.

### Phos-tag gel electrophoresis

Cortical neurons were infected with KD replacement viruses expressing GFP-Myc-tagged WT GluK2 or single point GluK2 mutants (A657T, M867I or S868A), all tagged at the N-terminus with GFP and Myc. Seven days post-infection, cells were treated with 1µM PMA for 20 min, lysed in 2× Laemmli sample buffer and subjected to 6% Phos-tag gel electrophoresis containing 50µM Phos-tag acrylamide (Fujifilm) to separate phosphorylated and non-phosphorylated protein species. Following electrophoresis, gels were washed twice with 10mM EDTA for 10 min each and once with transfer buffer prior to blotting. Membranes were probed with anti-myc antibody

### SUMOylation assay

HEK293 cells were co-transfected with 2µg of N-terminally GFP-Myc-tagged GluK2 constructs (WT, A657T, M867I, S868A, S868D or K886R), together with 0.1µg FLAG-Ubc9 and 0.2µg FLAG-SUMO1 to enhance SUMO conjugation. 24h post-transfection, cells were washed once with 1×PBS and lysed in buffer containing 20mM Tris-HCl (pH 7.4), 137mM NaCl, 1% Triton X-100, 20mM N-ethylmaleimide, 0.1% SDS, and cOmplete™ Protease Inhibitor Cocktail (Merck). Lysates were incubated on ice for 30 min and centrifuged at 13,500 rpm for 30 minutes at 4°C. The resulting supernatants were incubated with pre-washed GFP trap beads (ChromoTek) for 1 hour on a rotating wheel at 4 °C. Beads were washed three times with 600µl of washing buffer (20mM Tris-HCl, pH 7.4; 137mM NaCl; 1% Triton X-100) at 4000 rpm for 2 min, with 10 min incubations on a rotating wheel at 4°C between washes. After the final wash, 40µl of 2× Laemmli sample buffer was added to the beads, and samples were heated at 95°C for 10 min. Proteins were resolved by SDS-PAGE and analyzed by Western blotting using anti-FLAG and anti-GFP antibodies

### Acyl-PEGyl Exchange Gel Shift (APEGS) assay

To detect GluK2 palmitoylation, cortical neurons were infected at DIV14 with KD replacement viruses expressing GFP-Myc-tagged GluK2 WT, A657T or M867I. Seven days post-infection, palmitoylation was assessed using the APEGS assay, performed as previously described (Yucel *et al*. 2023; Yucel *et al*. 2024). Samples were subjected to SDS-PAGE and Western blotting, and membranes were probed with anti-Myc antibody.

### Endo H and PNGase F assay

Cortical neurons were infected at DIV12 with KD replacement viruses expressing GluK2 WT or A657T. Seven days post-infection, cells were washed with 1× PBS and lysed in buffer containing 20mM Tris-HCl (pH 7.4), 137mM NaCl, 1% Triton X-100 and protease inhibitors. Lysates were kept on ice for 30 minutes and centrifuged at 13,500 rpm for 20 minutes at 4 °C. Protein concentrations were determined using the BCA assay kit (Pierce, Thermo Scientific). 5µg of protein were treated with Endo H or PNGase F (New England Biolabs) according to the manufacturer’s instructions to assess protein trafficking between the ER and the Golgi apparatus.

### GluK2 degradation assays

Cortical neurons were infected with KD replacement viruses expressing GFP-Myc-tagged GluK2 WT, M867I or A657T. Seven days post-transduction, cells were treated 20µM MG132 (Sigma, Cat# 474790) or 20µM bafilomycin (Hello Bio, Cat# HB1125) for 0, 4 or 8 hours. Following treatment, cells were lysed and subjected to SDS–PAGE and Western blotting. Membranes were probed with anti-GluK2 antibody.

### Live surface staining and antibody feeding assay

These assays were carried out as described previously (Fletcher-Jones *et al*. 2024). Briefly, GFP-Myc-tagged GluK2 constructs (WT, M867I or A657T) were transfected into hippocampal neurons cultured on 25 mm coverslips. Seven days post-transfection, live neurons were cooled for 5 min to reduce internalisation and incubated with anti-GFP antibody in conditioned medium for 20 min at room temperature to label surface GluK2. Cells were washed three times with 1×PBS and fixed with pre-warmed 4% paraformaldehyde (PFA; Thermo Fisher) for 12 min. Fixation was quenched with 100mM glycine in PBS, followed by three additional washes with 1×PBS. After fixation, cells were permeabilised and blocked in 3% bovine serum albumin (BSA) with 0.1% Triton X-100 in PBS for 20 min. Surface GluK2 was visualised with Cy3-conjugated donkey anti-chicken IgY (1:400; Jackson ImmunoResearch, Cat# AB_2340363) in 3% BSA, while total GluK2 was detected using anti-GFP antibody followed by Alexa Fluor 488–conjugated donkey anti-chicken IgY (1:400; Jackson ImmunoResearch, Cat# AB_2337390).

To track GluK2 internalisation, hippocampal neurons plated on 25 mm coverslips were transfected with GFP-Myc-tagged GluK2 WT, M867I or A657T at DIV9–12. Seven days post-transfection, live cells were incubated with anti-GFP antibody for 45 minutes. Cells were then stimulated with 1µM PMA for 20 minutes. Following stimulation, surface-bound antibody was stripped by two quick washes with acidic PBS (pH 2.5). Cells were subsequently washed three times with cold 1× PBS, fixed with 4% PFA and permeabilised as described above. Internalised GluK2 was stained with Cy3-conjugated donkey anti-chicken IgY while total GluK2 was detected using anti-GFP antibody followed by Alexa Fluor 488–conjugated goat anti-chicken IgY.

### Ca^2+^ imaging

HEK293 cells were transfected with GFP-Myc-GluK2 WT(Q), WT(R), M867I, A657T or GFP together with R-GECO as a Ca²⁺ indicator (1:1 DNA ratio) in 25 mm glass bottom dishes. 48 hours post-transfection, conditioned medium was replaced with 2ml Tyrode’s buffer (25mM HEPES, 119mM NaCl, 2.4mM KCl, 2mM CaCl₂, 2mM MgCl₂ and 30mM glucose). Live-cell imaging was performed on a widefield microscope (Wolfson Bioimaging Facility, University of Bristol) using a 40× objective. The R-GECO channel was acquired with a 50 ms exposure time with 2×2 binning. Kainic acid (100 µM; Tocris, cat. #7065) was added at 0 min, and time-lapse images were acquired for 1800 frames at the minimum possible interval, for a total duration of 2 min.

Time-lapse image stacks were analysed in Fiji (ImageJ; https://fiji.sc/). For each cell, the region of interest (ROI) was drawn around the stimulated cell and fluorescence intensity over time was extracted using Plot Z-axis Profile. The resulting fluorescence trace (F) was exported and analysed in Excel and GraphPad Prism. Baseline fluorescence (F0) was calculated as the mean fluorescence during the first second of recording. Fluorescence values at each time point were normalised to baseline and expressed as F/F0.

### Acquisition and analysis for immunostaining

Confocal imaging was performed using a Leica SP8 confocal laser scanning microscope (Wolfson Bioimaging Facility, University of Bristol) with an HC PL APO CS2 63×/1.40 NA oil-immersion objective. All imaging settings were kept constant across conditions and across all independent replicates to allow quantitative comparisons. Z-stacks were collected with a 0.30 µm step size. For each condition, ≥10 cells per experiment were imaged from at least three independent dissections.

Images were analysed in Fiji. Z-stacks were maximum-intensity projected, and ROIs of approximately identical size were placed on three secondary dendrites per cell based on the total GluK2 signal. Mean fluorescence intensity was measured for each dendritic ROI and averaged to generate one value per cell. Surface GluK2 and internalised GluK2 signals were normalised to total GluK2 expression.

### Normalisation and statistical analysis

Surface expression was normalised to total expression, and total expression was normalised to GAPDH. Phosphorylated, SUMOylated and palmitoylated GluK2 signals were normalised to the corresponding non-modified GluK2 signals, respectively. For glycosylation analyses, Endo H-modified GluK2 was normalised to total GluK2 expression. For each independent experiment, normalised values for each condition were further scaled by dividing each value by the mean of all conditions within that experiment.

All graphs and statistical analyses were performed using GraphPad Prism (version 10). Statistical comparisons were performed using an unpaired t-test for two groups. For comparisons involving more than two groups, one-way or two-way ANOVA was used, followed by Tukey’s multiple-comparisons test. Data are presented as mean ± SEM. N indicates the number of independent replicates or the number of cells analysed from at least three independent dissections, as stated in the figure legends.

## Acknowledgements

The authors also gratefully acknowledge the Wolfson Bioimaging Facility (University of Bristol, UK) for their support and assistance in this work.

